# ATF3 Preserves Macrophage Cholesterol Homeostasis during Metabolic Stress

**DOI:** 10.64898/2026.11.03.764124

**Authors:** Amelia M. Pearson, Yusuf Barudi, Jihye Kang, Langley Williams, Charles N. Vallez, Cassidy J. Ulanowicz, Nandan S. Gokhale, Senad Divanovic, Ty D. Troutman

## Abstract

**Background:** Macrophages must coordinate immune activation with maintenance of cellular cholesterol homeostasis, yet the transcriptional mechanisms integrating these processes remain incompletely understood. Activating transcription factor 3 (ATF3) suppresses inflammatory and interferon responses and has been implicated in macrophage lipid metabolism, suggesting a role in coordinating immune and sterol-responsive transcriptional programs.

**Methods:** Wild-type and ATF3-deficient bone marrow-derived macrophages were analyzed by bulk RNA sequencing following Toll-like receptor stimulation and under conditions of sterol loading or depletion. Cholesterol content, cytokine production, and sterol-responsive gene expression were quantified. Human metabolic dysfunction-associated steatotic liver disease (MASLD) datasets and Kupffer cell-specific ATF3 knockout mice were used to assess disease relevance in vivo.

**Results:** ATF3 deficiency enhanced inflammatory and interferon-responsive transcriptional programs while suppressing a sterol-responsive gene module enriched for SREBP2 target genes. This phenotype was conserved across diverse Toll-like receptor stimuli and accompanied by persistent *Ch25h* upregulation. Despite exhibiting elevated cellular cholesterol, ATF3-deficient macrophages retained transcriptional responsiveness to sterol loading and depletion while demonstrating reduced induction of SREBP2-dependent genes across conditions. IFNAR blockade reduced interferon-responsive gene expression but did not normalize *Ch25h* or sterol-responsive gene expression. In contrast, CRISPR-mediated reduction of *Ch25h* partially restored sterol-responsive gene expression in inflammatory ATF3-deficient macrophages. In human MASLD, hepatic ATF3 expression positively correlated with disease severity. In mice, ATF3 was preferentially induced in Kupffer cells during metabolic stress, and Kupffer cell-specific deletion of ATF3 increased liver injury and Kupffer cell lipid accumulation.

**Conclusions:** ATF3 supports sterol-responsive transcriptional programs while restraining inflammatory activation in macrophages. ATF3 loss results in elevated cellular cholesterol, impaired SREBP2-dependent gene expression, and persistent derepression of *Ch25h*, which contributes to but does not fully explain the sterol phenotype. These findings identify ATF3 as a coordinator of macrophage adaptation to inflammatory and metabolic stress and support a protective role for Kupffer cell ATF3 during metabolic liver disease.

## Introduction

Macrophages are central regulators of lipid homeostasis and inflammation, two interconnected processes that drive the development of atherosclerosis and other chronic metabolic diseases^1–3^. In response to tissue injury, infection, or metabolic stress, macrophages must coordinate inflammatory activation with programs that maintain intracellular cholesterol/lipid balance. Failure to appropriately regulate these responses can result in excessive lipid accumulation, maladaptive inflammatory responses, and tissue damage^1–3^. Therefore, defining the mechanisms that integrate metabolic and inflammatory adaptation in macrophages is critical for understanding disease pathogenesis.

Cellular cholesterol homeostasis is maintained through opposing sterol-responsive transcriptional programs. When cellular cholesterol is low, sterol regulatory element-binding protein 2 (SREBP2) promotes cholesterol uptake and biosynthesis through induction of genes including *Ldlr* and enzymes of the mevalonate pathway^4^. Conversely, excess cholesterol activates liver X receptor (LXR)-dependent pathways that facilitate cholesterol efflux through targets including *Abca1*and *Apoe*^5^. Inflammatory signaling can directly influence these pathways through interferon-driven induction of cholesterol 25-hydroxylase (CH25H), which generates the oxysterol 25-hydroxycholesterol (25-HC), a potent inhibitor of SREBP2 activation^6^. Although inflammatory, interferon, and sterol-responsive pathways are extensively interconnected^1,3,7^, the transcriptional regulators that coordinate these responses during inflammatory stress are not fully understood.

Activating transcription factor 3 (ATF3) is a stress-responsive transcription factor^8^ that functions as a key negative regulator of inflammatory and type I interferon responses in macrophages^9–13^. In addition to its established immunomodulatory roles, ATF3 has been implicated in macrophage lipid metabolism through its regulation of foam-cell formation and protection from atherosclerosis^14,15^. Moreover, ATF3 expression and genomic occupancy increase in Kupffer cells during high fat diet-induced metabolic stress and integrative genomic analyses have identified ATF3 as a candidate regulator of disease-associated Kupffer cell transcriptional remodeling^16^. Together, these findings identify ATF3 as a potential regulator linking inflammatory signaling and sterol metabolism.

Here, we investigated whether ATF3 coordinates inflammatory and cholesterol-responsive transcriptional programs in macrophages. We identify ATF3 as a critical regulator of macrophage cholesterol homeostasis that supports SREBP2-dependent sterol-responsive gene expression across diverse inflammatory and metabolic conditions. Mechanistically, persistent derepression of *Ch25h* contributed to reduced sterol-response gene expression in inflammatory ATF3-deficient macrophages. Consistent with these findings, loss of ATF3 increases macrophage lipid accumulation in vitro and promoted Kupffer cell lipid accumulation and exacerbated liver injury during metabolic stress in vivo. Collectively, these findings establish ATF3 as a key transcriptional regulator linking inflammatory signaling to sterol homeostasis.

### Clinical Implications

Macrophage dysfunction contributes to the development of atherosclerosis, metabolic dysfunction-associated steatotic liver disease (MASLD), and other cardiometabolic diseases characterized by chronic inflammation and lipid accumulation. We identify ATF3 as a regulator that helps macrophages maintain cholesterol-responsive gene expression while limiting excessive inflammatory activation. Loss of ATF3 disrupted these adaptive responses, leading to cholesterol accumulation and heightened inflammatory signaling.

In a mouse model of metabolic liver disease, Kupffer cell-specific deletion of ATF3 increased Kupffer cell lipid accumulation and liver injury, supporting the importance of this pathway during chronic metabolic stress. These findings suggest that impaired macrophage adaptation to lipid and inflammatory stress may contribute to tissue injury and disease progression. A better understanding of how ATF3 maintains macrophage cholesterol homeostasis could reveal new therapeutic strategies to limit inflammation-driven tissue damage in atherosclerosis, MASLD, and related cardiometabolic disorders.

## Methods

### Ethics Statement

All animal studies were approved by Cincinnati Children’s Hospital Medical Center Institutional Animal Care and Use Committee at and follows institutional and internationally recognized animal welfare standards.

### Mice

Mice were maintained on a C57BL/6J background in the Troutman laboratory vivarium. The *Atf3* flox mice were provided by the Clifford Woolf laboratory^17,18^. The *Clec4f-Cre-tdTomato* mice are available from The Jackson Laboratory (Strain #:033296; RRID: IMSR_JAX:033296)^19^. Control (Atf3^fl/wt^, Clec4f-cre^+/-^) and Kupffer cell-specific ATF3-deficient (Atf3^fl/fl^, Clec4f-cre^+/-^) littermates were generated for in vivo studies. Experiments were performed using male littermates at starting ages of 6-9 weeks of age. Lyz2-Cre mice were obtained from The Jackson Laboratory (Strain#: 031674; RRID: IMSR_JAX:031674)^20^. Control (Atf3^fl/fl^, Lyz2-cre^-/-^) and myeloid-specific ATF3-deficient (Atf3^fl/fl^, Lyz2-cre^+/-^) littermates were used to generate bone marrow-derived macrophages (BMDMs).

### Model Diets

Mice were fed either a MASLD-model diet (Research Diets, D09100310i; 40 kcal% Fat [Mostly Palm Oil], 20 kcal% Fructose and 2% Cholesterol) for up to 55 weeks or the discontinued Amylin liver NASH diet (Research Diets, D09100301; 40 kcal% Fat [Mostly Primex], 20 kcal% Fructose and 2% Cholesterol) for up to 30 weeks. Body conditioning score^21^ and body weight were recorded weekly. Blood samples were collected at baseline and every 4 weeks thereafter. At study termination, serum was collected for downstream analysis.

### Cell culture

Preparation of BMDMs was previously described^19^ and further detailed in the supplemental methods.

### In Vitro Conditions

BMDMs were stimulated with lipopolysaccharide (LPS) (100 ng/mL; Invivogen), Pam3CSK4 (100 ng/mL; Invivogen), Pam2CSK4 (100 ng/mL; Invivogen), CpG DNA (100 nM; Invivogen), or imiquimod/R837 (1 µg/mL; Invivogen) for up to 24 h. Foam cell formation was induced by treatment with oxLDL (100 µg; Invitrogen, L34357) for 24 h. Lipid deprivation (LD) was achieved by culturing cells in medium containing 1% lipid-depleted FBS (Kalen Biomedical, 880100). Where indicated, cells were pretreated for 1 h with anti-IFNAR (10 µg/mL; BioXcell, BE0241) prior to experimental stimulation, and antibody was replenished during media change.

### Crispr-Cas9

Gene deletion was performed on bone marrow cells as described^22^ and detailed in the supplemental methods.

### Quantitative real-time PCR

Total RNA was extracted from BMDMs using Monarch Spin RNA Isolation Kit (Monarch #T2110L). RNA concentration was measured by Qubit fluorometer (Invitrogen, 14175095) before reverse transcription with Superscript III RT (Invitrogen 18080085). Quantitative real-time PCR (qPCR) was performed on a QuantStudio 7 Flex Real-Time PCR System (Life Technologies) and analyzed using the comparative Ct method and normalized to *Hprt*. Primers were validated by melt curve analysis and amplification efficiency. Primer sequences are listed in Table S1.

### ELISA

IL-6 and TNF-α concentrations were quantified by sandwich ELISA. Details are in the supplemental methods.

### ISGF3 Reporter Assay

L929 cells stably expressing an ISGF3-responsive secreted *Gaussia* luciferase reporter were generated by lentiviral transduction using the pTRIP ISGF3-Gluc transfer plasmid (Gift from Ram Savan)^23^. Viral particles were packaged in HEK-293T cells together with psPAX2 (Addgene) and the VSV-G envelope from pMD2.G (Addgene) and concentrated using Takara Lenti-X concentrator. Transduced L929 cells were single cell cloned by limiting dilution, and an IFN-responsive clone was selected for subsequent experiments.

Reporter cells were maintained in RPMI containing 5% FBS and 1% pen/strep/glutamine. Cells were seeded in 96-well plates (50,000/well) and incubated at 37°C. After 24 h, media was replaced with conditioned medium from BMDM cultures. Following incubation, reporter cell supernatants were collected and clarified by centrifugation (800 x g, 8 minutes, 4°C). *Gaussia* luciferase activity was measured immediately or after storage at -80°C using the *Gaussia* Luciferase Glow Assay (Thermo Cat#16160). Supernatants were added to coelenterazine-containing assay buffer in opaque 96-well plates (Thermo Cat#15042), and luminescence was quantified using a Synergy H1 plate reader.

### Total Cholesterol Measurement

BMDMs were lysed in lipid lysis buffer (50 mM Tris-HCl, 150 mM NaCl, 0.1% Triton X-100) and collected by scraping before being transferred to microcentrifuge tubes. Lysates were centrifuged (15,000 x g, 15 min) to remove cellular debris. Total cholesterol levels were quantified using the Amplex Red Cholesterol Assay Kit (Invitrogen, A12216) according to the manufacturer’s instructions. Assays were performed in black-walled microplates (Invitrogen, M33089) to minimize fluorescence spillover.

### Poly A RNA-seq

Sequencing libraries were prepared as described^24,25^ and are detailed in the supplemental methods.

### Sequencing Data Analysis

FASTQ files were mapped to the mouse reference genome assembly GRCm39 using STAR with default parameters^26^. Alignment quality was assessed using FASTQC, and alignments were visually inspected in the UCSC Genome Browser and/or Integrated Genomics Viewer. Aligned reads were converted into HOMER tag directories for downstream analysis^27^. Expression counts were generated using HOMER’s ‘analyzeRepeats’ script with the options - count exons, - condenseGenes, - noadj, producing a table of raw exon-associated read counts per gene for each experiment. Differential expression analysis was performed using raw counts by DESeq2^28^. Differentially expressed genes were defined as genes with a Benjamini-Hochberg adjusted p-value < 0.05 and an absolute fold change > 2, corresponding to an absolute log2 fold change > 1. For visualization by principal-component analysis, heatmaps, and selected gene expression plots, raw counts were transformed using regularized log transformation with DESeq2.

Gene set enrichment analysis was performed using DESeq2 Wald statistics from each contrast as ranked inputs. Ranked gene lists were tested against Hallmark gene sets, and enrichment was summarized by normalized enrichment score (NES) and Benjamini-Hochberg adjusted p-value. Sterol and interferon-stimulated gene modules were derived from the Reactome cholesterol biosynthesis and interferon alpha/beta signaling gene sets (MSigDB C2:CP:REACTOME), respectively, with curated additions and removals (detailed in Supplemental Table 2). The NF-κB module was assembled from established NF-κB-dependent inflammatory mediators, effector enzymes, and feedback regulators. All modules were restricted to genes with baseMean > 100 across samples prior to scoring. DESeq2 rlog-transformed expression values were centered for each gene using the mean expression of ATF3 WT samples at 0 hours (untreated condition) and scaled by the pooled within group standard deviation across genotype-by-time groups. Module scores were calculated as the mean gene-level z-score for all retained genes within each module. Scores were plotted across the time course by genotype and ligand.

### Serum Analyte Analysis

Whole blood was collected into EDTA-coated tubes (BD, 365974) and centrifuged (10,000 x g, 10 min). Serum was aliquoted and stored at -80°C until analysis.

In house measurement of serum alanine aminotransferase (ALT) was performed using the Catachem ALT kit (Catachem, C164-0A) following manufacturer’s instructions. Comprehensive serum chemistry analysis was performed by IDEXX BioAnalytics (Test Code 60514).

Serum chemistry data were compiled from multiple male cohorts of GAN-fed mice at harvest. The ALT/AST dataset included 48 KC-ATF3 WT mice and 44 KC-ATF3 KO mice spanning a body weight range of 29 – 57 grams and 9 – 45 weeks on diet and included both fasted and non-fasted animals. Associations between body weight and serum analytes were assessed using linear regression with genotype, body weight, and their interaction as fixed effects. Cohort and fasting status were included as covariates.

In an independent experiment, 13 KC-WT and 10 KC-ATF3 KO male mice were fed the GAN diet for ∼45 weeks. Animal mass was tracked every two weeks, and serum was collected at weeks 0, 30 and at study conclusion. Serum ALT levels were assessed via the Catachem ALT kit. Data were statistically analyzed using a linear mixed-effects model with genotype, time, the genotype by time interaction, and mass as fixed effects, and mouse as a random intercept (R: lme4). At each timepoint, the genotype difference was estimated as the difference in estimated marginal (least-squares) means between KC-ATF3 KO and KC-WT mice at equal body mass and P values were Šidák-adjusted for the three timepoints.

### Spectral Flow Cytometry of Non-Parenchymal Cells

In brief, non-parenchymal cells were stained with a viability dye (Live/Dead NIR, Invitrogen) and anti-CD16/CD32 to block Fc receptors, then labeled with fluorescently conjugated primary antibodies. Flow cytometry data were acquired using a Cytek Aurora. Antibodies used are listed in the Major Resources Table.

### Preparation of Non-Parenchymal Cells

Mice were euthanized by isoflurane overdose and exsanguinated prior to isolation of liver non-parenchymal cells. Livers were retrograde perfused for 2 minutes at 3 mL/minute with Ca^2+^/Mg^2+^ free HBSS (Gibco, 14175095) supplemented with 0.5mM EGTA (bioWORLD, 40121266-3), 0.5 mM EDTA (Invitrogen, 15575020), and 20 mM HEPES (TEKNOVA, H1090), followed by perfusion with 40 mL digestion buffer consisting of HBSS containing Ca^2+^/Mg^2+^ (Gibco, 14025134)) supplemented with 20 mg/mL collagenase, 20 µg/mL DNase1, and 20 mM HEPES. Livers were removed, minced, and further digested for 20 minutes at 37°C with gentle rotation in fresh digestion buffer. To minimize digestion-induced transcriptional changes, perfusion and digestion buffers contained 1 µM flavopiridol (Sigma, F3055).

After digestion, hepatocytes were removed by two low-speed centrifugation steps (50 x g, 2 minutes) and the remaining cell suspension was filtered through a 70-micron strainer. Non-parenchymal cells were enriched by centrifugation in 20% isotonic percoll (Sigma, P4937) (600 x g in 50 mL, 15 min, 4 °C, followed by density gradient separation (28% Optiprep (Sigma, D1556), 1,400 x g, 25 min, 4°C, no break, slow acceleration). Cells recovered from the interface were subjected to erythrocyte lysis, washed, and resuspended in PBS.

### Measurement of Neutral Lipids in Liver Macrophages

Non-parenchymal cells were isolated as described above and stained with 2 µM bodipy (Invitrogen, D3922) in PBS for 30 minutes at 37°C. Following PBS, washes, cells were stained for spectral flow cytometry as described above.

### Analyses of Public Data Sets

Human bulk liver RNA-seq in the form of annotated and normalized count matrix available on NCBI was from GSE135251^29^. Data was log_2_ transformed and analyzed for an association between hepatic ATF3 expression and NAS using Kendall’s rank correlation across the available 216 samples. P-values were adjusted across all genes using Benjamini-Hochberg correction.

## Results

### ATF3 represses inflammatory and interferon responses while promoting sterol-responsive gene expression during macrophage activation

Given that ATF3 regulates macrophage immune responses^9–13^ and has also been implicated in limiting foam-cell formation through transcriptional repression of the CH25H-25HC pathway^14,15^, we sought to define how ATF3 shapes transcriptional programs during macrophage activation. To address this question, we generated wild-type (WT) and ATF3-deficient BMDMs and profiled their transcriptional response across a Toll-like receptor 4 (TLR4) stimulation time course using lipopolysaccharide (LPS) treatment and bulk RNA sequencing (6 independent samples per genotype across 2 independent experiments). LPS effects dominated gene expression variance along PC1 and PC2 (**Fig. 1A**). Across the time course, 908 unique genes were differentially expressed (DESeq2: absolute log_2_ fold-change > 1and adjusted p-value < 0.01) between genotypes, including 264 at homeostasis (**Fig. 1B**). Genotype-dependent differences had non-overlapping patterns across timepoints, with many uniquely detected at either 6 hours or 24 hours of LPS treatment (**Fig. S1A).** This result may indicate that ATF3 shapes distinct transcriptional programs in macrophages at early versus late stages of an LPS response. Consistent with previous studies identifying ATF3 as a negative regulator of immune signaling^9,10,12,13^, the expected induction of inflammatory and interferon-associated programs was observed in ATF3-deficient macrophages.

**Figure 1.**
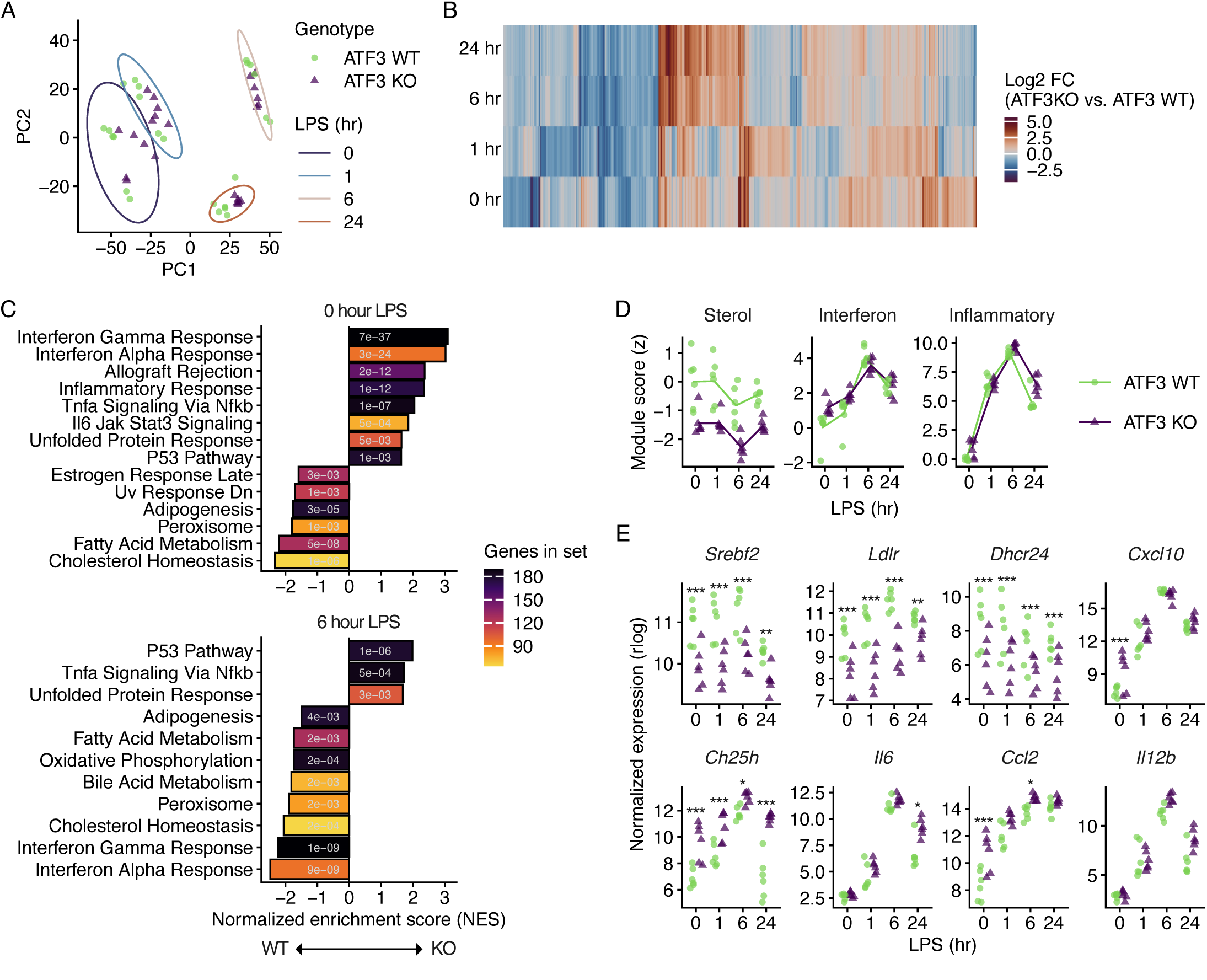
ATF3 loss derepresses inflammatory and interferon programs and suppresses sterol biosynthesis in LPS-stimulated macrophages. BMDMs from ATF3-WT and ATF3-KO mice were stimulated with LPS and harvested at 0, 1, 6, and 24 h for RNA-seq (n = 6/genotype/timepoint across two independent experiments with batch included in the model). (**A**) Principal component analysis of regularized-log (rlog) transformed expression. Ellipses denote genotype-pooled 95% probability regions per timepoint (multivariate t-distribution fit). (**B**) Hierarchical clustering of log₂ fold change values (ATF3-KO vs. ATF3-WT) for 908 differentially expressed genes (adjusted P < 0.01 and |log₂FC| > 1) at the indicated timepoints (DESeq2, Wald test, Benjamini–Hochberg correction). (**C**) Gene set enrichment analysis (GSEA, Hallmark collection) of the KO-vs-WT contrast at the indicated timepoint. Bars show normalized enrichment scores (NES) with positive NES values indicating enrichment in KO and negative values in WT. Bar fill denotes gene set size and inset values inflammatory programs across the LPS time course. Scores are the mean of per-gene z-scored rlog expression across each gene set. (**E**) Representative expression (rlog) of indicated genes across the LPS time course. Points are individual biological replicates.

To determine whether loss of ATF3 altered coordinated transcriptional responses to LPS, we next performed gene set enrichment analysis (GSEA) using Hallmark gene sets and DESeq2 Wald statistic ranked gene lists comparing ATF3-deficient and WT macrophages at each timepoint. At baseline, ATF3 deficiency was associated with enrichment of Hallmark pathways including Interferon Gamma Response, Interferon Alpha Response, Inflammatory Response, and TNF-α Signaling via NF-κB (**Fig. 1C**). In contrast, the Cholesterol Homeostasis gene set was negatively enriched (**Fig. 1C**), indicating lower expression of this transcriptional program in ATF3-deficient macrophages relative to WT cells. Following LPS stimulation, the interferon-associated genes sets were no longer differentially enriched between genotypes, whereas the inflammatory response pathways remained positively enriched and the cholesterol homeostasis program remained negatively enriched (**Fig. 1C**; **Fig. S1B**).

To visualize temporal changes in the enrichment of these gene sets, we constructed gene modules exemplifying ATF3-dependent sterol-response (sterol), interferon-response (interferon), and NF-κb–dependent inflammatory transcription (inflammatory) in macrophages. A module score for each sample was calculated as the mean z-scored expression across the constituent genes in reference to the ATF3 WT macrophages at homeostasis (0 h LPS). The interferon module was elevated in ATF3-deficient macrophages at baseline and converged on WT macrophages upon LPS stimulation, whereas the inflammatory module remained elevated (**Fig. 1D, E**). Conversely, the sterol module was significantly reduced in ATF3-deficient macrophages across the entire LPS time course, exemplified by *Srebf2*, *Ldlr*, and *Dhcr24* (**Fig. 1D, E**). Among the most strongly derepressed genes in ATF3-deficient macrophages across the time course was *Ch25h*, which encodes CH25H and converts cholesterol to 25-HC. In turn, 25-HC inhibits activation of a SREBP2, a key transcriptional regulator of cholesterol biosynthesis^6^.

Together, these findings supported the established role of ATF3 in restraining inflammatory^9,10^ and type I interferon responses^12,13^ and provided temporal resolution to this regulation. In parallel, we identified a previously underappreciated role for ATF3 in sustaining the expression of sterol-responsive genes at homeostasis and under LPS challenge. Unlike the interferon program, this sterol-suppressive signature was present at homeostasis and persisted through the acute and late phases of the LPS response. The coincident derepression of *Ch25h* alongside suppression of the sterol module suggested a potential link between these phenotypes in ATF3-deficient macrophages.

### ATF3-dependent regulation of sterol, interferon, and inflammatory programs is conserved across toll-like receptor stimuli

The analyses above identified suppression of sterol-response genes as a prominent feature of ATF3-deficient macrophages during LPS stimulation. However, because TLR4 signal transduction proceeds through both MyD88- and TRIF-dependent pathways, it was unclear whether reduced sterol gene expression was specific to TLR4 or represented a broader feature of ATF3-dependent regulation of the TLR family. This was relevant because in addition to distinct ligand specificities, TLRs also differ in the cellular compartmentalization from which they initiate signaling, which could influence response dependencies. We therefore tested whether ATF3-dependent regulation of inflammatory, interferon, and sterol-responsive transcriptional programs was conserved across TLR activation. WT and ATF3-deficient BMDMs were stimulated with ligands for TLR1/2 (Pam3CSK4), TLR2/6 (Pam2CSK4), TLR9 (CpG DNA), TLR7 (imiquimod/R837), and TLR4 (LPS).

ATF3-deficient macrophages secreted more TNF-α after 6h and more IL-6 after 24h when stimulated through TLR4, TLR7, or TLR9 (**Fig. 2A, B**). In contrast, neither cytokine differed between genotypes following cell-surface and MyD88-dependent TLR1/2 with Pam3CSK4. This result indicated a potential ligand specificity in how ATF3 might regulate expression of certain NF-κB targets. To determine whether ATF3 potentially regulates the secretion of type I interferons, we transferred conditioned media from macrophages to cultured L929 fibroblasts expressing an ISGF3-driven *Gaussia* luciferase reporter. Conditioned media from unstimulated ATF3-deficient macrophages induced greater ISGF3 activity than WT conditioned media, consistent with elevated basal production of type I interferons (**Fig. 2C**). ISGF3 reporter activity was also elevated in ATF3-deficient macrophage media following 6h stimulation with CpG or R837, but not at 24h. Despite the absence of detectable genotype-dependent differences in TNF-α/IL-6 secretion, conditioned media from ATF3-deficient macrophages following stimulation of TLR1/2 with Pam3CSK4 or TLR2/6 with Pam2CSK4 induced greater ISGF3 reporter activity. In contrast, the elevated ISGF3 reporter activity observed at baseline was not sustained upon LPS stimulation, consistent with expression of the ISG module (**Fig. 1D**). Together, these findings indicate that ATF3 restrains both inflammatory and interferon output across multiple TLR stimuli.

**Figure 2.**
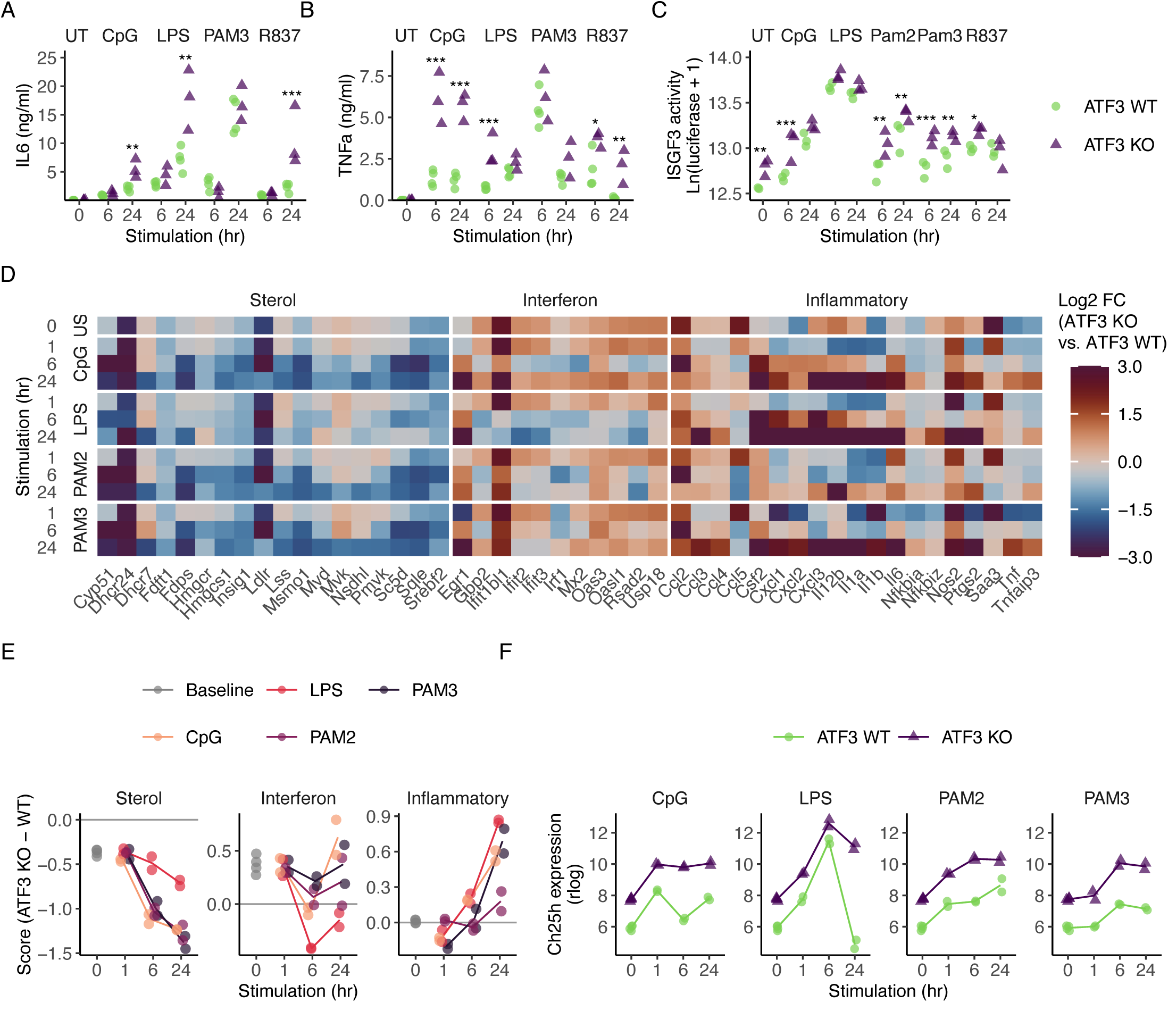
ATF3 restrains inflammatory and interferon output and suppresses sterol biosynthesis across multiple TLR ligands. BMDMs from ATF3-WT and ATF3-KO mice were stimulated with the indicated TLR ligands and analyzed by ELISA for (**A**) IL6 or (**B**) TNFα secretion (n = 3-4/genotype/ligand; representative of 2 independent experiments). Points are individual replicates. (**C**) ISGF3 reporter activity following treatment with conditioned media from stimulated BMDMs (n = 3/genotype/ligand from one experiment). Asterisks denote genotype differences based on estimated marginal means with Benjamini-Hochberg correction (*P < 0.05, **P < 0.01, ***P < 0.001, ****P < 0.0001). (**D**) Log₂ fold change (ATF3-KO vs. ATF3-WT) for differentially expressed genes (adjusted P < 0.01 and |log₂FC| > 1; DESeq2, Wald test, Benjamini–Hochberg correction) in the sterol, interferon, and inflammatory modules (n = 2/genotype/ligand from one experiment). (**E**) Module scores differences (ATF3 KO – ATF3 WT) for the sterol, interferon, and inflammatory programs. Scores represent the mean of per-gene z-scored rlog expression values across each module. The common 0 h baseline is displayed as the starting point for each stimulus trajectory (gray). (**F**) *Ch25h* expression (rlog) across stimulation time course. Points are individual biological replicates; shared unstimulated samples are plotted at 0 h in each facet.

To determine whether ATF3 regulates distinct downstream transcriptional programs through different TLRs, we conducted a time course on macrophages treated with CpG, LPS, Pam2CSK4, or Pam3CSK4 and performed RNA-seq (n = 2 per condition). Across this time course, 1,554 unique genes (DESeq2: absolute log_2_ fold-change > 1 and adjusted p-value < 0.01) were differentially expressed between genotypes, ranging from a sum of 1,642 DEGs across LPS stimulated cells to 993 DEGs across Pam2CSK4 stimulated cells. Next, we compared the relative expression of individual genes in the sterol, interferon, and inflammatory response modules defined here (**Fig. 2D**) and found that individual members of the modules had some ligand specific expression patterns when comparing TLR1/2, TLR2/6, and TLR9 stimulated macrophages, but generally followed the patterns observed for TNF-α/IL-6 secretion and ISGF3 reporter activity (**Figs. 2A-C**). Similar trends were observed when comparing module score differences between ATF3-deficient versus ATF3-sufficient macrophages (**Fig. 2E**). Notably, TLR4 stimulated ATF3-deficient macrophages again showed an acute de-repression (1 h) in interferon stimulated gene expression that became suppressed beneath wild-type levels at later time points (6 and 24 h) (**Fig. 1C, D and Fig. 2D, E**). Whereas inflammatory and interferon modules displayed time- and ligand-specific differences, the sterol module was consistently reduced across every ligand and time point in ATF3-deficient macrophages (**Fig. 2D-E**).

Because suppression of the sterol program was conserved across ligands, we next examined *Ch25h*, one of the most consistently derepressed genes in ATF3-deficient macrophages (**Fig. 1E**). *Ch25h* expression was elevated in ATF3-deficient macrophages across all ligands and timepoints examined and remained elevated at 24 h post-stimulation even when expression in WT macrophages had returned to the baseline (**Fig. 2F**). Given the established role of 25HC in regulating cholesterol metabolism, the persistent derepression of *Ch25h* represented a potential link between ATF3 deficiency and suppression of sterol-responsive gene expression. We therefore next examined whether the altered sterol program in ATF3-deficient macrophages was accompanied by changes in cholesterol homeostasis.

### ATF3-deficient macrophages exhibit elevated cholesterol despite intact cholesterol sensing

Macrophage ATF3 has been shown to limit foam cell formation and protect against atherosclerosis^14^; however, its role under lipid-rich, inflammatory conditions remains poorly defined. To model these conditions, WT and ATF3-deficient macrophages were loaded with oxLDL for 24 hours before LPS stimulation (**Fig. 3A**). Under basal conditions, ATF3-deficient macrophages contained significantly higher total cholesterol than WT cells in both unloaded and oxLDL-loaded states. Following LPS stimulation, cholesterol levels increased in WT macrophages, eliminating the genotype-dependent difference (**Fig. 3B**). The observation that ATF3-deficient macrophages contained more cholesterol at homeostasis prompted us to investigate whether this was accompanied by changes in cholesterol-responsive transcriptional programs using RNA-seq (n = 7-9 per condition across three independent experiments). Genes exhibiting significant genotype-dependent expression (p-adjust < 0.05 in at least one condition) were intersected with a curated Hallmark and Gene Ontology cholesterol-associated gene sets (Supplemental Table 2) and organized into functional pathways within cholesterol homeostasis (**Fig. 3C**).

**Figure 3.**
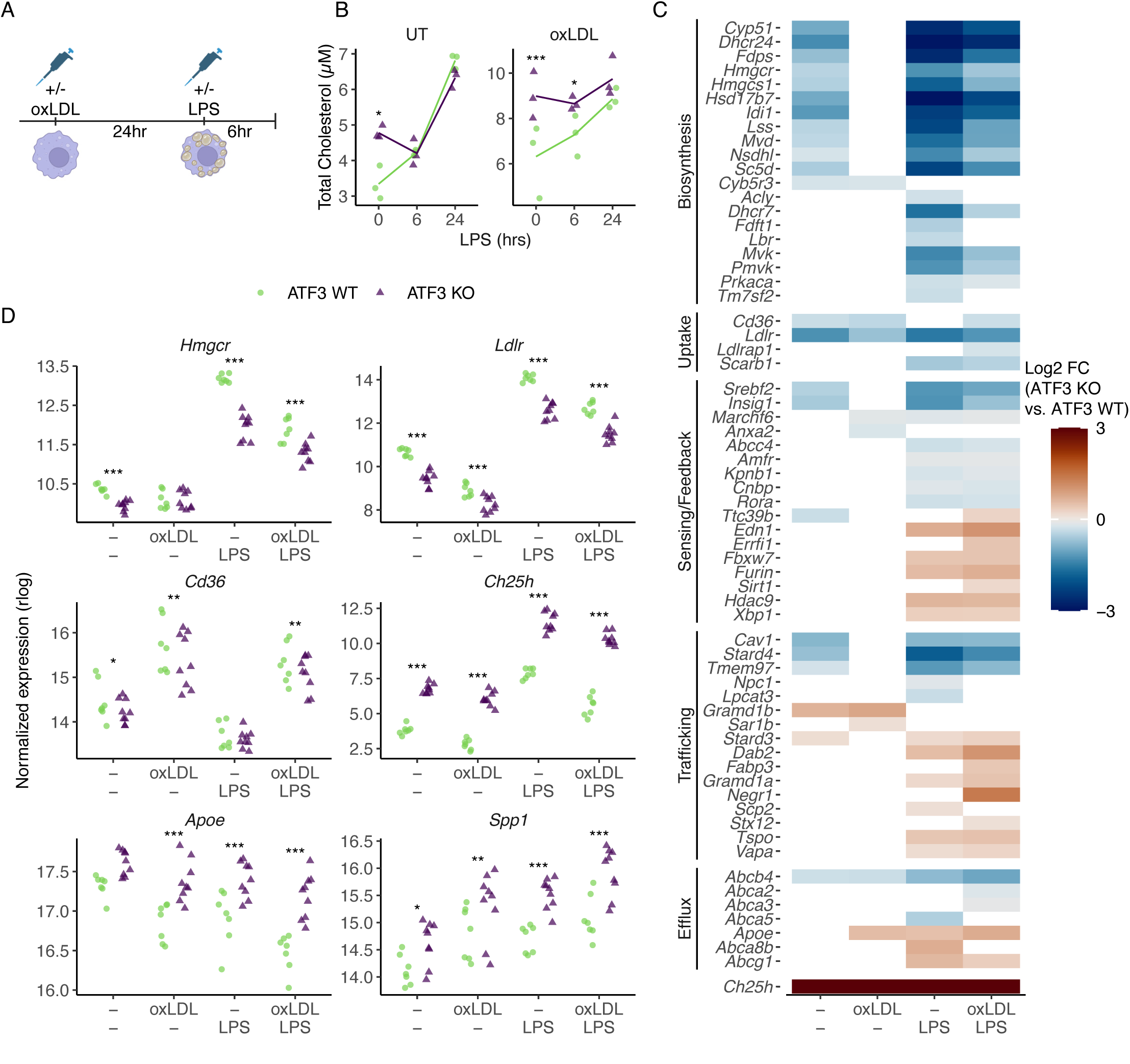
ATF3 supports cholesterol homeostasis during inflammatory foam cell formation. (**A**) Experimental schematic. BMDMs from ATF3-WT and ATF3-KO mice were loaded with oxLDL for 24 h prior to stimulation with LPS. (**B**) Total cellular cholesterol content measured by Amplex Red assay (n = 3/genotype/timepoint, representative of two experiments). Points are individual biological replicates. Statistical significance was assessed by mixed-effects modeling, followed by ANOVA and multiple-comparison-adjusted pairwise testing. (*P < 0.05, **P < 0.01, ***P < 0.001). (**C**) Log₂ fold change (ATF3-KO vs. ATF3-WT) for differentially expressed genes (adjusted P < 0.05; DESeq2, Wald test, Benjamini–Hochberg correction) grouped by cholesterol-related function. (**D**) Expression (rlog) of indicated genes across treatment conditions. Points represent individual biological replicates (n = 7-9/genotype/condition across 3 independent experiments with batch included in the model).

Largely, ATF3 loss was associated with broad suppression of genes involved in cholesterol acquisition. Canonical cholesterol biosynthesis genes, including *Hmgcr*, were consistently expressed at lower levels in ATF3-deficient macrophages, with genotype differences becoming more pronounced following LPS stimulation (**Fig. 3C-D**). Similarly, expression of *Ldlr* and *Cd36* was reduced in ATF3-deficient macrophages, indicating coordinated suppression of cholesterol synthesis and uptake programs (**Fig. 3C-D**). Notably, these transcriptional differences were modest under basal conditions but were amplified following inflammatory activation.

In contrast to the reduced expression of cholesterol synthesis and uptake pathways, *Ch25h*, *Apoe*, and *Spp1* were consistently increased in ATF3-deficient macrophages (**Fig. 3C-D**). OxLDL loading reduced the number of genotype-dependent differences across cholesterol-related pathways, whereas LPS stimulation elicited broad transcriptional differences regardless of lipid-loading status (**Fig. 3C-D**). Collectively, elevated cholesterol content coincided with reduced expression of cholesterol acquisition pathways and altered expression of multiple cholesterol-associated pathways in ATF3-deficient macrophages (**Fig. 3B-D**). Because SREBP2 is a central regulator these processes, we next investigated whether altered SREBP2-dependent gene regulation contributes to this phenotype.

### Impaired sterol-program activation in ATF3-deficient macrophages is independent of inflammatory activation

The sterol-responsive transcriptional program was initially identified in the context of inflammatory stimulation, raising the possibility that reduced expression of cholesterol-responsive genes was secondary to basal inflammatory activation rather than a direct defect in sterol regulation. To distinguish between these possibilities, WT and ATF3-deficient macrophages were subjected to lipid deprivation (LD), a treatment condition established to induce SREBP2 activation in the absence of exogenous inflammatory cues^4^. LD expectedly increased expression of SREBP2 target genes *Hmgcr*, *Insig1*, and *Ldlr* in WT macrophages (**Fig. 4A**), confirming activation of the canonical sterol-deprivation response. Although ATF3-deficient macrophages retained the ability to induce these genes, the rate of induction was markedly reduced relative to WT cells (**Fig. 4A**). In contrast, *Ch25h* expression remained consistently elevated in ATF3-deficient macrophages throughout the time course (**Fig. 4A**). These findings indicate that impaired activation of the sterol-response program in ATF3-deficient macrophages is not dependent on inflammatory stimulation and persists under conditions that normally activate SREBP2. As persistent derepression of *Ch25h* across basal, inflammatory, and lipid-deprived conditions identified it as a candidate mediator of the sterol-suppressed phenotype (**Fig. 1E, 2F, 3D, 4A**), we next sought to determine what maintains elevated *Ch25h* expression in ATF3-deficient macrophages.

**Figure 4.**
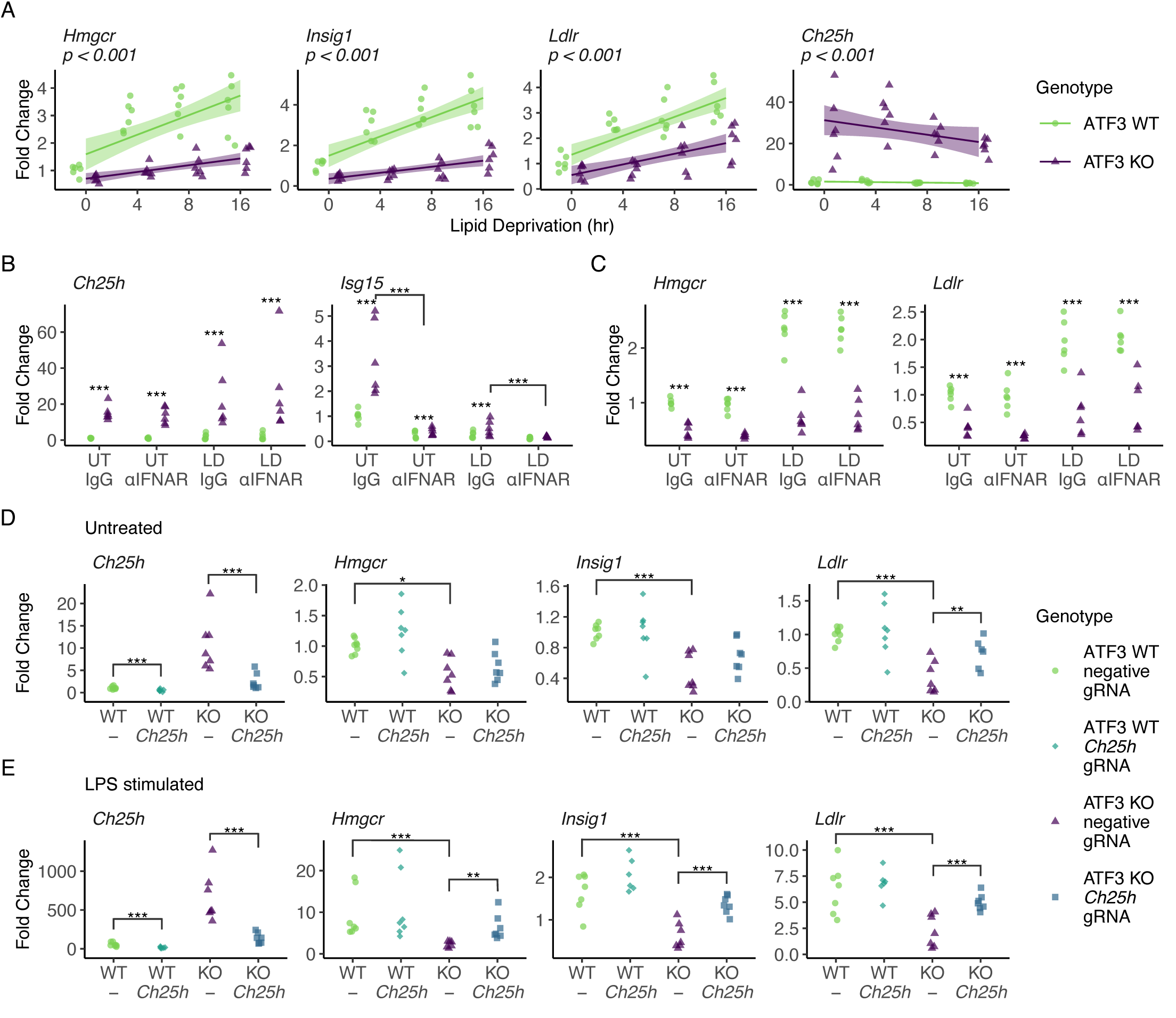
ATF3 supports SREBP2-dependent gene expression, partially through repression of *Ch25h*. BMDMs from ATF3-WT and ATF3-KO mice were subjected to LD and analyzed by qPCR. (**A**) Relative expression of *Hmgcr*, *Insig1*, *Ldlr*, and *Ch25h* during LD (n = 6/genotype/time point across two independent experiments with batch included in the model). (**B-C**) BMDMs were pre-treated with an anti-IFNAR antibody and harvested after 8 h LD. Relative expression of *Ch25h* and *Isg15* (**B**) and *Hmgcr* and *Ldlr* (**C**) (n = 6/genotype/condition across two independent experiments). (**D-E**) Relative expression of *Ch25h*, *Hmgcr*, *Insig1*, and *Ldlr* in control-and Ch25h-targeted CRISPR BMDMs under basal (D) or LPS-stimulation (E) conditions (n = 6-7/genotype/condition across three independent experiments). Points represent individual biological replicates. Statistical significance was assessed by linear or mixed-effects modeling, followed by ANOVA and multiple-comparison-adjusted pairwise testing. (*P < 0.05, **P < 0.01, ***P < 0.001).

Because *Ch25h* is a well-characterized interferon-stimulated gene^6^ and ATF3-deficient macrophages exhibit elevated secretion of type 1 interferons and expression of interferon stimulated genes at homeostasis^12^ (**Figs. 1C, 1D, 2C, 2D, 2E**), we asked whether persistent *Ch25h* expression was maintained by autocrine type I interferon signaling. To test this possibility, WT and ATF3-deficient macrophages were treated with an IFNAR-blocking antibody under basal or lipid-deprived conditions. Under treatment with an isotype control antibody, *Isg15* expression was expectedly higher in ATF3-deficient macrophages and was substantially reduced following IFNAR blockade, largely diminishing the genotype-associated difference (**Fig. 4B**). In contrast, *Ch25h* expression remained elevated in ATF3-deficient macrophages after IFNAR blockade under both untreated and lipid-deprived conditions (**Fig. 4B**). Likewise, suppression of *Hmgcr* and *Ldlr* expression persisted despite IFNAR blockade (**Fig. 4C**).

Although these findings excluded ongoing IFNAR signaling as the primary driver of the sterol phenotype, they did not address whether elevated *Ch25h* expression directly contributes to the impaired activation of sterol-response genes. We therefore tested whether reducing *Ch25h* expression could restore activation of the sterol program in ATF3-deficient macrophages. To accomplish this, WT and ATF3-deficient bone marrow cells were transduced with CRISPR/Cas9 constructs targeting *Ch25h* prior to differentiation into macrophages. Consistent with our previous observations, ATF3-deficient macrophages displayed significantly lower expression of sterol-response genes under both basal and inflammatory conditions compared to WT controls. *Ch25h* reduction partially restored expression of sterol response gene expression in LPS-stimulated ATF3-deficient macrophages (**Fig. 4E**). However, *Ch25h* reduction did not significantly increase sterol gene expression in lipid-deprived ATF3-deficient macrophages (**Fig. S2A**).

Collectively, these findings establish ATF3 as a regulator of macrophage adaptation to sterol and inflammatory stress. Loss of ATF3 resulted in exaggerated inflammatory responses with impaired activation of sterol-responsive gene programs, phenotypes that were maintained under lipid-rich conditions and only partially explained by derepression of *Ch25h*. We next sought to determine whether ATF3-dependent regulation of macrophage cholesterol homeostasis and inflammatory activation is relevant in a disease setting characterized by chronic lipid accumulation and inflammation.

### ATF3 expression in Kupffer cells limits liver injury in diet-induced obesity

Metabolic dysfunction-associated steatohepatitis (MASH) is characterized by lipid accumulation and chronic inflammation^30^, conditions that parallel the environmental cues used in our in vitro studies. In humans, hepatic ATF3 expression has been associated with worsened disease severity^31^. To evaluate this association, we analyzed public bulk liver RNA sequencing from 216 patients spanning the spectrum of MASLD and confirmed a positive correlation between ATF3 expression and histological disease scores (GSE135251) (**Fig. 5A**)^29^. Although elevated ATF3 expression is associated with disease progression, murine genetic studies suggest that this elevation is not a disease driver but instead a protective response because ATF3 loss in multiple tissues exacerbates MASLD pathology^31–34^. Moreover, ATF3 expression in Lyz2^+^ cells (e.g. monocytes, some macrophages, some granulocytes) has been implicated in limiting disease progression^35^. These findings suggest that macrophage-associated ATF3 may broadly contribute to protection from metabolic liver injury.

**Figure 5.**
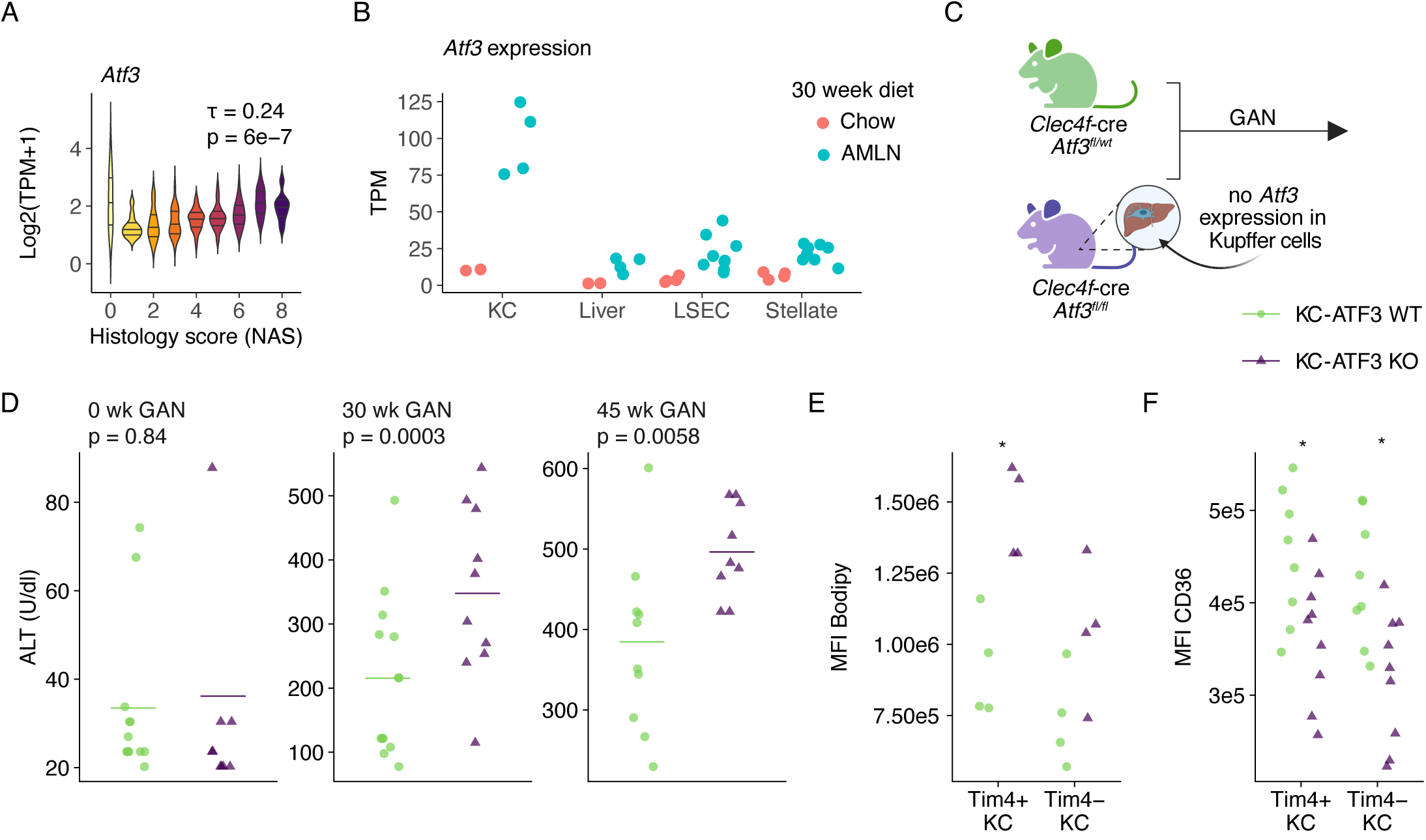
Kupffer cell ATF3 limits liver injury during diet-induced obesity. (**A**) Kendall correlation between NAS scores and *Atf3* expression in human liver RNA-seq datasets (GSE135251). (**B**) *Atf3* expression in whole liver, liver sinusoidal endothelial cells (LSEC), stellate cells, and Kupffer cells from WT mice maintained on chow or AMLN diet for 30 weeks (n = 2-4/chow and n = 4-9/AMLN). (**C**) Generation of KC-ATF3 WT (*Atf3^fl^*^/wt^; *Clec4f*-Cre^+/-^) and KC-ATF3 KO (*Atf3^fl^*^/fl^; *Clec4f*-Cre^+/-^) mice. (**D**) KC-ATF3 KO mice and KC-ATF3 WT littermate controls were maintained on GAN diet for up to 50 weeks and serum ALT was measured at the indicated diet timepoints (n = 10/genotype from one independent experiment). Statistical significance was assessed by linear mixed effects modeling and time point specific differences determined with estimated marginal means. (**E-F**) Neutral lipid accumulation (n = 4/genotype representative of 2 independent experiments) (**E**) and CD36 surface expression (n = 8-9/genotype across two independent experiments) (**F**) in Clec4f^+^Tim4^+^ and Clec4f^+^ Tim4^-^Kupffer cells after 40 weeks of GAN feeding, assessed by spectral flow cytometry. Statistical significance was determined by Student’s t test (*P < 0.05, **P < 0.01, ***P < 0.001). Points represent individual biological replicates.

Because bulk liver transcriptomics primarily reflects hepatocyte-driven transcripts, we next examined hepatic cell populations from WT mice following 30 weeks of high fat diet (AMLN) feeding. Among hepatic cell populations, *Atf3* induction was most pronounced in Kupffer cells (**Fig. 5B**). As liver resident macrophages, Kupffer cells are continuously exposed to lipid and inflammatory stresses and play a central role in coordinating local inflammatory and metabolic responses during MASLD progression^36,37^. Consistent with their relevance to the phenotypes identified in our in vitro studies, our previous multi-omic analysis of Kupffer cells during AMLN feeding identified ATF3 as a predicted regulator of MASLD-induced transcriptional remodeling and demonstrated increased ATF3 expression and genomic occupancy during diet-induced metabolic stress^16^. Based on these observations, we next tested the consequences of Kupffer cell-specific ATF3 deletion during MASLD.

To test this hypothesis, we generated Kupffer cell-specific ATF3 knockout (KC-ATF3 KO; *Atf3^fl/fl^; Clec4f-Cre^+/-^*) mice and WT (KC-WT; *Atf3^fl/wt^; Clec4f-Cre^+/-^*) littermate controls. Mice were maintained on a Gubra Amlyn NASH diet^38^ (GAN) for up to 50 weeks (**Fig. 5C**). We assessed the relationship between mouse weight and measures of peripheral liver damage, serum alanine transaminase (ALT) and aspartate aminotransferase (AST), across independent GAN-cohorts of varying lengths of time. Although weight gain over time did not different between genotypes, KC-ATF3 KO mice had approximately 7.5 U/L per gram body weight serum ALT activity and 8.9 U/L per gram AST activity relative to KC-WT mice (**Fig. S3A-C**). These findings suggested that loss of Kupffer cell ATF3 increased susceptibility to liver injury as metabolic burden increased. Consistent with this interpretation, an independent cohort maintained on GAN for 47 weeks showed that ALT did not differ between genotypes at baseline but was elevated in KC-ATF3 KO mice after 30 and 47 weeks of GAN feeding (**Fig. 5D**).

As loss of ATF3 resulted in altered lipid homeostasis in vitro, we quantified neutral lipid accumulation in Kupffer cells. Tim4^+^ KC-ATF3 KO Kupffer cells accumulated significantly more neutral lipid than KC-WT controls, as assessed by BODIPY staining (**Fig. 5E**). Furthermore, both Tim4^+^ and Tim4^-^KC-ATF3 KO Kupffer cells exhibited reduced surface CD36 expression (**Fig. 5F**). These findings are consistent with our in vitro observations that ATF3-deficient macrophages accumulate excess cholesterol while exhibiting reduced expression of cholesterol acquisition pathways, including *Cd36* (**Fig. 3B, D**). Despite these alterations in lipid content, bulk RNA-sequencing of KC-WT and KC-ATF3 KO Kupffer cells isolated after 30 weeks GAN diet identified relatively few differentially expressed genes (**Fig. S4A-C**). This observation parallels our in vitro studies, where prolonged exogenous lipid loading reduced genotype-dependent differences in cholesterol-responsive transcriptional programs (**Fig. 3C-D**). Together, these findings demonstrate that loss of ATF3 promotes lipid accumulation within Kupffer cells and is associated with increased markers of liver injury during chronic metabolic stress.

## Discussion

Here, we identify a role for ATF3 in the maintenance of macrophage sterol homeostasis. ATF3-deficient BMDMs accumulated cholesterol and had impaired expression of cholesterol biosynthesis and uptake genes at homeostasis, during inflammatory activation, and under lipid-deprived conditions. Notably, these defects persisted despite neutralization of type I interferon signaling, separating ATF3-dependent control of sterol metabolism from its established role in suppressing interferon-responsive gene expression. In vivo, selective deletion of ATF3 in Clec4f^+^ Kupffer cells led to increased liver injury during experimental MASLD and altered lipid handling in Kupffer cells. Together, our results support a model by which ATF3 coordinates macrophage cholesterol homeostasis through a distinct mechanism from the canonical type I interferon/CH25H regulatory pathway.

One literature-supported interpretation of the sterol phenotype would be that it results as a downstream consequence of ATF3 constraint of interferon signaling and interferon-induced expression of *Ch25h*. ATF3 directly represses *Ch25h* promoter acetylation and expression^14^, and *Ch25h* is often described as an interferon-stimulated gene whose product 25-HC potently suppresses SREBP2 activation^6^. However, our results do not fully support this model. Suppression of the sterol module was present at homeostasis and across treatment with multiple TLR ligands, whereas the interferon module was both ligand- and time-dependent, and no longer elevated upon TLR4 stimulation with LPS. Further, blockade of type I interferon signaling suppressed *Isg15* to near wildtype levels but had minimal impact on genotype-dependent differences in expression of *Ch25h*, *Hmgcr*, and *Ldlr*. This result reproduced observations made in Labzin et al, which also shows that *Ch25h* remains derepressed in ATF3-deficient macrophages independently of ongoing type I interferon signaling, in contrast to *Isg15*. CRISPR-mediated Ch25h reduction partially restored sterol-response gene expression in LPS-stimulated but not lipid-deprived ATF3-deficient macrophages, consistent with the 25-HC restraint of SREBP2 being most evident during inflammatory activation. Therefore, ATF3 supports sterol-responsive transcription through a mechanism that is distinct from autocrine type I interferon signaling and only partially explained by *Ch25h*-dependent feedback.

Several limitations should be considered in the interpretation of our results. Although ATF3-deficient macrophage accumulated cholesterol and had suppressed expression of sterol-responsive genes, the molecular basis of these phenotypes remains unresolved. Reduced expression of SREBP2 target genes could reflect impaired redistribution of sterols to the ER, which cannot be resolved by our approaches. Alternatively, because ATF3 functions as a transcription factor, impaired expression of sterol-responsive genes may arise downstream of sterol sensing in the ER. Likewise, incomplete knockdown of *Ch25h* limits conclusions regarding the contribution of 25HC to SCAP/INSIG mediated suppression of SREBP2 with the observed phenotypes. Finally, despite reproducible effects of Kupffer cell-specific ATF3 deletion on liver injury and Kupffer cell lipid accumulation, transcriptomic analyses of purified Kupffer cells and whole liver tissue failed to identify corresponding genotype-dependent transcriptional programs. These findings raise the possibility that genotype-dependent phenotypes effects occur within cellular states that are not readily captured by bulk sequencing approaches; or that post-transcriptional, metabolic, or other undefined mechanisms contribute to the in vivo phenotype. Future studies directly examining ER sterol pools, SREBP2 activation, and ATF3-dependent transcriptional regulation will be required to establish causal mechanisms linking ATF3 to macrophage cholesterol homeostasis and liver protection during MASLD.

Our in vivo studies extend these observations to chronic metabolic disease. ATF3 expression was strongly induced in Kupffer cells during diet-induced metabolic stress, consistent with previous evidence identifying ATF3 as a stress-responsive factor in tissue macrophages^16^. Interestingly, ATF3 activity in macrophages protected against hepatic steatosis by enhancing fatty acid oxidation through transcriptional regulation of FOXO1^35^. These experiments used deletion in Lyz2^+^ macrophages and orthogonal overexpression in adoptively transplanted bone-marrow derived macrophages but could not resolve causal macrophage subsets. Our experiments here provide some resolution as we found that specific loss of ATF3 in Clec4f^+^ Kupffer cells increased macrophage lipid accumulation and was associated with heightened markers of liver injury during prolonged dietary challenge. Notably, these phenotypes were observed despite limited transcriptional differences at late stages of disease. Similar convergence was observed in vitro following prolonged lipid loading, where transcriptional differences became less pronounced despite persistent differences in cholesterol content, suggesting that lipid exposure may reduce ATF3-dependent transcriptional differences while preserving underlying defects in cellular cholesterol homeostasis. Together with our findings, these observations support a broader role for ATF3 in coordinating macrophage lipid metabolism during chronic lipid stress. Importantly, the mechanism identified here may not fully explain the in vivo phenotype. *Ch25h* expression was substantially lower in Kupffer cells than in BMDMs, suggesting that *Ch25h*-dependent regulation may contribute less prominently in tissue-resident liver macrophages. In this context, the FOXO1-dependent regulation of fatty acid oxidation described by Hu et al. and the incomplete rescue achieved by *Ch25h* knockdown both support ATF3 coordination of multiple metabolic programs in macrophages.

Together, these studies identify ATF3 as a coordinator of macrophage inflammatory and sterol-responsive programs. Loss of ATF3 exaggerates inflammatory activation while simultaneously impairing maintenance of a sterol-responsive transcriptional network, resulting in cholesterol accumulation despite apparent compensatory suppression of cholesterol acquisition pathways. Persistent derepression of *Ch25h* contributes to this phenotype but does not fully explain it. In vivo, Kupffer-cell ATF3 limits lipid accumulation and protects against liver injury during chronic metabolic stress. Collectively, these findings support a model in which ATF3 promotes macrophage adaptation to inflammatory and lipid stress by maintaining appropriate sterol-responsive transcriptional responses.

## Non-standard Abbreviations and Acronyms

LD: lipid deprivation
oxLDL: oxidized low-density lipoprotein
BMDM: bone marrow derived macrophage

## Acknowledgements

This work was made possible using the Cincinnati Children’s Hospital Medical Center Research Flow Cytometry Facility (RRID:SCR_022635), Integrated Pathology Research Facility (RRID:SCR_022637), Genomics Sequencing Facility (RRID:SCR_022630), and the Cincinnati Children’s Data Management and Analysis Center (RRID:SCR_022625). *Atf3* flox mice were generously provided by the Clifford Woolf laboratory at the Boston Children’s Hospital.

## Author Contributions

Conceptualization, AMP, TDT, CNV, NSG, CP, and SD; Methodology, AMP, TDT, YB, LW, and CJU; Formal Analysis, AMP and TDT; Investigation, AMP, YB, JK, TDT, LW, and CJU; Data Curation, TDT; Writing – Original Draft, AMP and TDT; Writing – Review and Editing, AMP, JK, and TDT; Funding Acquisition, TDT.

## Supplemental Material

Supplemental Methods

Figure S1-4

Table S1-2

Major Resources Table

---

**Supplemental Figure 1. AT**F3 **shapes distinct transcriptional programs in macrophages at early versus late stages of an LPS response**. BMDMs from ATF3-WT and ATF3-KO mice were stimulated with LPS and harvested at 0, 1, 6, and 24 h for RNA-seq (n = 6/genotype/timepoint across two independent experiments with batch included in the model). (**A**) UpSet plot displaying the size of individual and intersecting gene sets identified. The horizontal bars on the left represent the total number of differentially expressed genes (DEGs) per contrast. The vertical bars on the top show the size of each intersecting subset, defined by the connected black circles in the matrix below. (**B**) Gene set enrichment analysis (GSEA, Hallmark collection) of the KO-vs-WT contrast at the indicated timepoint. Bars show normalized enrichment scores (NES) with positive NES values indicating enrichment in KO and negative values in WT. Bar fill denotes gene set size and inset values inflammatory programs across the LPS time course.

**Supplemental Figure 2. Ch25h reduction partially rescues SREBP2 target gene expression during lipid deprivation.** (**A**) Relative expression of *Ch25h*, *Hmgcr*, *Insig1*, and *Ldlr* in control- and *Ch25h*-targeted CRISPR BMDMs under 8h LD conditions (n = 6-7/genotype/condition across three independent experiments). Points represent individual biological replicates. Statistical significance was assessed by linear modeling, followed by ANOVA and multiple-comparison-adjusted pairwise testing. (*P < 0.05, **P < 0.01, ***P < 0.001).

**Supplemental Figure 3. Kupffer cell ATF3 protects against liver damage beyond a body-weight threshold during diet-induced obesity.** (**A–B**) Relationship between body weight and serum ALT (**A**) and AST (**B**) in GAN-fed KC-ATF3 WT and KC-ATF3 KO mice. Associations were analyzed by multiple linear regression. (**C**) Body weight of KC-WT and KC-ATF3 KO mice throughout the HFD time course. Statistical significance was assessed by linear modeling, followed by ANOVA.

**Supplemental Figure 4. Transcriptional differences between liver macrophages. (A)** Principal component analysis bulk RNA-seq from FACS purified Ly6CHi macrophages (Mo-Mac) or Tim4+Clec4f-tdTomato+ Kupffer cells (KC) from the mice of the indicated genotypes (n = 4 from one experiment). Data are from DESeq2 regularized log transform expression data. **(B)** MA plot comparing DESeq2-identified differentially expressed genes (adjusted p value < 0.05, absolute(log_2_ fold-change) > 1) between cell types from ATF3 WT mice. **(C**) Representative expression (rlog) of indicated genes. Points are individual biological replicates.

## Supplemental Methods

### Generation of bone marrow derived macrophages

Bone marrow-derived macrophages (BMDMs) were generated from *Atf3^fl/fl^, LyzM-cre^+/-^* and *Atf3^fl/fl^, LyzM-cre^-/-^* mice. In brief, red blood cell lysed (eBioscience, 00-4333-57) bone marrow cells was isolated were seeded onto non-tissue culture treated petri dishes (BD cat. no. 351029) and differentiated for 4 days in RPMI supplemented with 10% fetal-bovine serum (FBS) (Corning, 10-040-CV), 1% pen/strep/glutamine (Cytiva, SV300821), 51µM 2-mercaptoethanol, and 20 ng/mL MCSF (Biolegend, 576406). MCSF was supplemented on day 3 of differentiation to a final concentration of 20 ng/mL. On day four, cells were washed once with 2mM EDTA (Invitrogen, 15575020) then incubated in 2mM EDTA at 4°C for 20-40 minutes to facilitate detachment. Macrophages were the collected by scraping and replated for use in experiments.

### Crispr-Cas9

Gene deletion was performed on bone marrow cells as previously described^40^. Briefly, single guide RNAs (sgRNA; 0.3nmol each, Synthego) were complexed with Cas9 nuclease (0.6µl, Integrated DNA Technologies) in nuclease free water for 10 minutes and electroporated into ten million red blood cell lysed bone marrow cells using a Lonza 4D-Nucleofector system (Lonza, AAF-1003X; setting mouse mature DC P3; pulse DN107). Following electroporation, cells were recovered in pre-warmed macrophage differentiation media at 37°C for 10 minutes, washed, and cultured under BMDM differentiation conditions. sgRNA sequences are provided in Table S1.

### ELISA

IL-6 and TNFα concentrations were quantified by sandwich ELISA. Plates were coated with capture antibody overnight at 4℃ blocked with 1% BSA in PBS for 2-3 hours at 37°C and washed 3 times with PBS-Tween. Diluted samples and standards were added and incubated overnight at 4℃ Following 3 washes, detection antibody was added for 2-3 hours at 37°C. Plates were washed 4 times, incubated with streptavidin-HRP (1:1000, Jackson Immunoresearch, 016-030-084) for 30 minutes at 37°C and washed 5 times with PBS-Tween. Color development was performed using OPD substrate (1 tablet/10mL; Sigma Aldrich P8412) in citrate Phosphate Buffer (0.5M Na2HPO4, 1M Citric Acid, 0.03% H2O2). Reactions were stopped with 1M H2SO4 and absorbance was measured at 490nm.

### Poly A RNA-seq

Briefly, RNA was isolated from cultured cells using RNA lysis buffer (0.110 M Tris-HCl pH 7.5, 0.55 M LiCl, 11 mM EDTA, 1.1% SDS, 5.5% DTT) and concentration was measured by Qubit fluorometer. 100ng RNA per sample was suspended in 50µL Ultrapure H2O (Invitrogen, AM9937), stored at -80°C, and utilized for subsequent library preparation.

To prepare sequencing libraries, mRNAs were enriched by incubation with Oligo d(T) Magnetic Beads (NEB, S1419S) and then fragmented/eluted by incubation at 94°C for 9 min. Poly A enriched mRNA was fragmented, in 2x Superscript III first-strand buffer with 10mM DTT (Invitrogen), by incubation at 94°C for 9 min, then immediately chilled on ice before the next step. The 10 μL of fragmented mRNA, 0.5 μL of Random primer (Invitrogen), 0.5 μL of Oligo dT primer (Invitrogen), 0.5 μL of SUPERase-In (Ambion), 1 μL of dNTPs (10 mM) and 1 μL of DTT (10 mM) were heated at 50°C for three minutes. At the end of incubation, 5.8 μL of water, 1 μL of DTT (100 mM), 0.1 μL Actinomycin D (2 μg/μL), 0.2 μL of 1% Tween-20 (Sigma) and 0.5 μL of Superscript III (Invitrogen) were added and incubated in a PCR machine using the following conditions: 25°C for 10 min, 50°C for 50 min, and a 4°C hold. The product was then purified with RNAClean XP beads according to manufacturer’s instruction and eluted with 10 μL nuclease-free water. The RNA/cDNA double-stranded hybrid was then added to 1.5 μL of Blue Buffer (Enzymatics), 1.1 μL of dUTP mix (10 mM dATP, dCTP, dGTP and 20 mM dUTP), 0.2 μL of RNase H (5 U/μL), 1.05 μL of water, 1 μL of DNA polymerase I (Enzymatics) and 0.15 μL of 1% Tween-20. The mixture was incubated at 16°C for 1 h and held at 4°C overnight. The resulting dUTP-marked dsDNA was purified using 28 μL of Sera-Mag Speedbeads (Thermo Fisher Scientific), diluted with 20% PEG8000, 2.5M NaCl to final of 13% PEG, eluted with 40 μL EB buffer (10 mM Tris-Cl, pH 8.5). The purified dsDNA (40 μL) underwent end repair by blunting, A-tailing and adaptor ligation using barcoded adapters (NextFlex, BioScientific). Libraries were PCR-amplified for 9 cycles, quantified using a Qubit dsDNA HS Assay Kit (Thermo Fisher Scientific) and sequenced on a NovaSeqX Plus (Illumina, San Diego, CA) in the CCHMC Genomics Sequencing Facility according to the manufacturer’s instructions.

